# bioRxiv: the preprint server for biology

**DOI:** 10.1101/833400

**Authors:** Richard Sever, Samantha Hindle, Ted Roeder, Sol Fereres, Olaya Fernández Gayol, Sanchari Ghosh, Martina Proietti Onori, Emma Croushore, Kevin-John Black, Linda Sussman, Janet Argentine, Wayne Manos, Marisol Muñoz, Josh Sinanan, Tracy K. Teal, John R. Inglis

## Abstract

The traditional publication process delays dissemination of new research, often by months, sometimes by years. Preprint servers decouple dissemination of research papers from their evaluation and certification by journals, allowing researchers to share work immediately, receive feedback from a much larger audience, and provide evidence of productivity long before formal publication. Launched in 2013 as a non-profit community service, the bioRxiv server brought preprinting to the life sciences and recently posted its 310,000th manuscript. The server now receives around ten million views per month and hosts papers spanning all areas of biology. Initially dominated by evolutionary biology, genetics/genomics and computational biology, bioRxiv has been increasingly populated by papers in neuroscience, cell and developmental biology, and many other fields. bioRxiv and its sister server, medRxiv, also played a critical role during the pandemic, rapidly disseminating new discoveries in immunology, virology and epidemiology related to the SARS-CoV-2 virus and its effects. Changes in journal and funder policies that encourage preprint posting helped drive adoption, as did the development of bioRxiv technologies that allow authors to transfer papers easily between bioRxiv and journals. A recent user survey found that 30% of authors post their preprints weeks to months before submitting to a journal whereas 55% post around the time of journal submission. Authors are motivated by a desire to share work early; they value the feedback they receive and very rarely experience any negative consequences of preprint posting. Rapid dissemination via bioRxiv is also encouraging new initiatives that experiment with the peer review process and the development of novel approaches to literature filtering and assessment.

## Introduction

Dissemination of scientific manuscripts has traditionally occurred only after the research has been formally evaluated by scientific journals (Sever, 2023). In the print era, the high marginal costs associated with distribution favored this coupling of evaluation and dissemination; only manuscripts that passed a certain bar set by the journal were published and incurred printing costs. The resulting delays to dissemination have often prompted scientists to share draft manuscripts informally among close colleagues, and more organized mechanisms for sharing preprints widely were piloted as early as the 1960s (Cobb, 2017).

Over the years, concerns about delayed dissemination have become more acute. The routine requirement among journals for external peer review became universal only in the past few decades (Sever, 2023), and authors increasingly feel that the demands made by reviewers and editors are lengthening the publication process still further (Vale, 2015). Moreover, the timescale of journal publication, which can take months to years (Royle, 2015), is increasingly at odds with the timescales on which scientists, in particular early career researchers, must demonstrate productivity when evaluated for appointments, tenure and grants (Sarabipour et al., 2019).

The advent of the Web offered an opportunity to decouple the dissemination of papers from their subsequent evaluation and certification by journals. The costs of dissemination online are significantly lower, reducing the financial argument for disseminating only peer-reviewed papers; online dissemination is almost immediate; and anyone with an Internet connection can view the work. The arXiv preprint server, launched in 1991 and currently hosted by Cornell University, has demonstrated the effectiveness of this approach (Ginsparg, 2011). Researchers in physics, computational science, mathematics, and various other disciplines routinely post manuscripts on arXiv prior to peer review, and by late 2025 the site had posted around 2.9M papers. Several attempts were made to replicate the approach in the biological sciences (Marshall, 1999; Nature Publishing Group, 2012; Rawlinson and Bloom, 2019). These were unsuccessful in part because of opposition from traditional publishers but also because there was little interest among biologists. More recently, however, the increasing pace of research, increasing dissatisfaction with delays caused by peer review, restricted availability of many published papers, and a general growth in enthusiasm for more openness and transparency in science communication have refocused attention on the potential for preprints in biology. bioRxiv was launched in 2013 in the hope that rapid sharing of biology preprints would eliminate delays to dissemination (Sever, 2026; Kaiser, 2013) and in doing so increase the pace of research itself (Quake, 2019). The purpose of this article is to summarize bioRxiv’s progress and potential and provide a general reference for the project.

## The launch of bioRxiv

bioRxiv was an initiative of Cold Spring Harbor Laboratory (CSHL), a non-profit research institute with a unique international reputation as both a leading research institute and a hub for scientific communication. CSHL has been a meeting place for scientists for more than 100 years and a center of professional scientific education for more than 50 years. The annual CSHL Symposium was central to the birth of molecular biology and genomics, and conferences at CSHL continue to attract thousands of scientists every year. The laboratory also has significant publishing expertise, as the originator of classic books and manuals and several academic journals published by its publishing division CSHL Press. It was therefore a natural steward for a community preprint server for the life sciences, and the initiative received strong encouragement from the laboratory’s leadership. bioRxiv was launched in 2013 following discussions with members of the academic community, librarians, and arXiv (Sever, 2026). Notably, following consultations with representatives of arXiv, the project was named “bioRxiv”, not “bio-arXiv”, to reduce the likelihood that users would mistakenly contact arXiv staff for bioRxiv technical support. In 2025, management of bioRxiv and its sister server medRxiv passed to openRxiv, a non-profit organization set up to ensure the servers’ long-term stability and independent governance by members of the scientific community.

### Technical basis for bioRxiv

Given the potentially vast number of biology preprints — several hundred thousand papers each year — it was clear that bioRxiv would require an industrial scale architecture that could process and display a high volume of submissions and stably accommodate millions of online readers with minimal downtimes. bioRxiv’s hosting and manuscript management sites would have to include state-of-the-art features biologists had come to expect of online journals and be able to accommodate both existing and future integrations with other participants in the scholarly communication ecosystem (e.g. search engines, indexing services, journals, and manuscript submission systems). After defining the specifications required, we partnered with Highwire Press, a company developed within Stanford University that is now owned by MPS Limited and has a proven record of more than 20 years in online manuscript hosting and technology development for clients including CSHL Press and the *British Medical Journal* (BMJ).

The submission side of bioRxiv is based on a BenchPress submission system adapted for preprint handling and automated transfer to the display site. The display side is based on modified Highwire JCore technology. Additional customization by CSHL developers uses JavaScript and external databases to enhance and supplement the display on the site and provide additional feeds and services. In addition, the site is integrated with the third-party Disqus and Hypothesis commenting/annotation tools. A significant difference from traditional journals is that the architecture needs to accommodate the ability to upload revised versions of papers at any time (Fig. 1). All preprints are assigned a single digital object identifier (DOI). Since 2019 the date of author approval has been embedded within the DOI suffix so the date of the original version of the article is easily discernable from a citation. Each version of the preprint receives a unique URL, with the DOI for the preprint defaulting to the most recent version of the paper posted (see below). Articles can be cited by DOI or version-specific URL identifier.

**Fig. 1.**
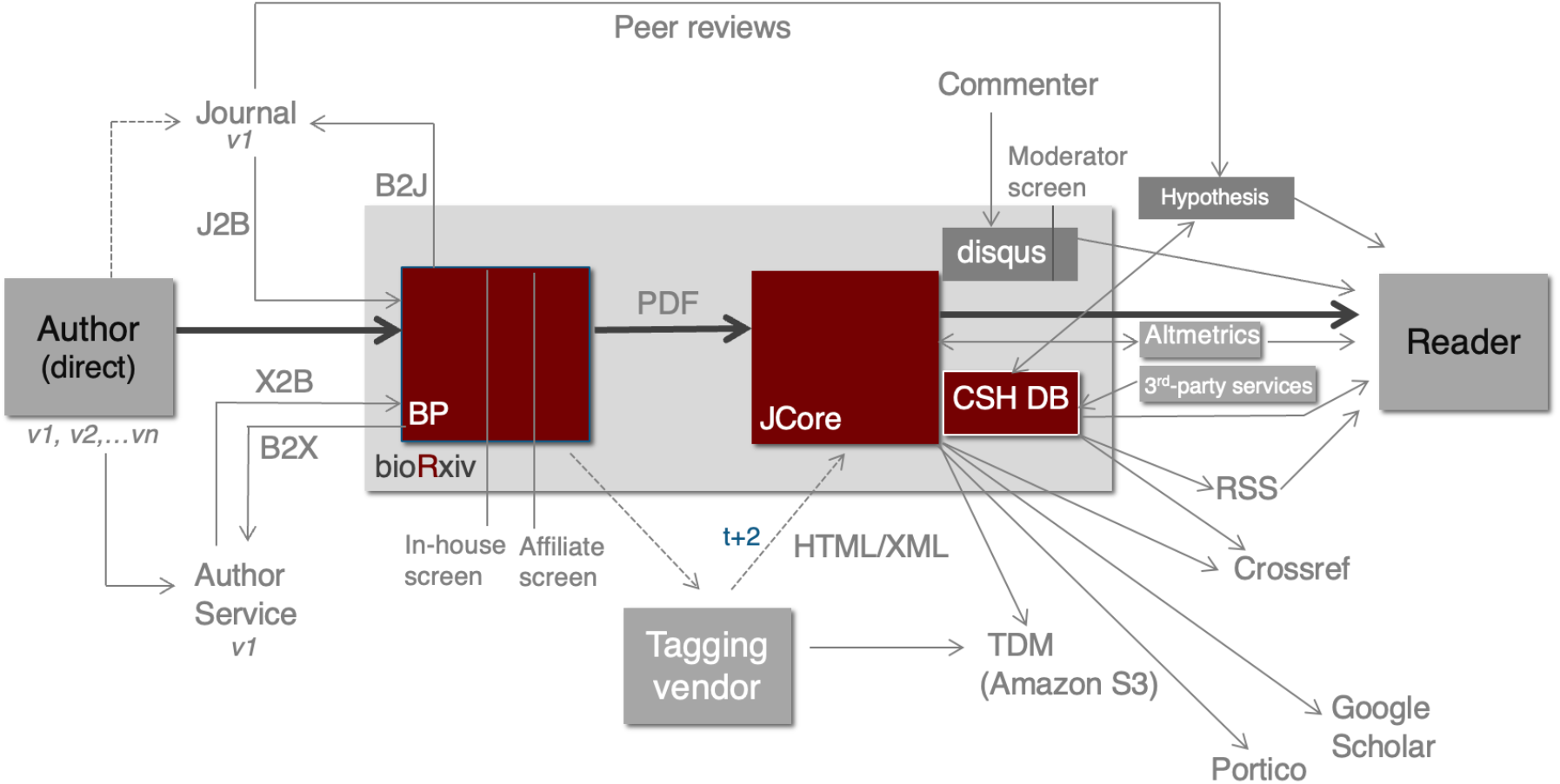
bioRxiv workflow. Authors submit papers through a BenchPress (BP) module either directly or via J2B from journals and other services. Manuscripts are then displayed publicly via the Highwire JCore module. CSHL databases (CSH DB) augment display and pass information to third parties. Papers are screened in BP and can also be sent to journals via B2J. HTML and XML are generated by a compositor. TDM, text and data mining repository.

bioRxiv is committed to permanency of the content posted. All content is therefore also deposited with the archiving service Portico, a not-for-profit organization committed to long-term preservation of scholarly material.

### Preprint screening

A defining feature of bioRxiv and medRxiv is that they do not perform peer review. Nevertheless, there is a need to screen papers to minimize the chance of posting of inappropriate material and maximize the content’s utility to readers. The bioRxiv screening process acts as a coarse filter for non-scientific/pseudoscientific content, non-biological/biomedical content, and potentially harmful content, as well as manuscripts solely comprising isolated data elements, and non-research articles such as recipes, textbook excerpts, narrative reviews and speculative theory. The decision to decline articles other than research papers, no matter how worthy, was a pragmatic one aimed at maximizing screening efficiency. It reduces subjectivity in screening and recognizes the reality that it is research rather than review/didactic content that suffers the distribution delays bioRxiv is intended to address (Sever, 2019).

bioRxiv screening is a two-stage process performed in a highly customized BenchPress environment. Papers first undergo an internal screen by bioRxiv staff, which includes automated plagiarism checks using Similarity Check software and search engines, as well as manual checks for spam and clearly inappropriate or incomplete content. Submissions are then further screened by a distributed group of bioRxiv Affiliates, all of whom are experienced scientists. This ensures that every article posted on bioRxiv has been viewed by a scientist. It is important to emphasize, however, that the screening process is a coarse, quick filter intended to minimize the likelihood that readers will encounter content that is not *bona fide* biological research. It does not guarantee or certify the content in any way, and readers must use their own judgment in assessing its validity as science.

Initially bioRxiv was intentionally restricted to basic biology: any clinical work was excluded. This restriction was partially lifted in 2015/2016 with the introduction of a pilot in which clinical research could be posted in two specific areas: epidemiology and registered clinical trials. Such papers had a specific screening process involving a group of medically qualified bioRxiv Clinical Affiliates. In 2019, the success of this pilot resulted in the launch of a dedicated preprint server for clinical research, medRxiv (Rawlinson and Bloom, 2019), and the bioRxiv Epidemiology and Clinical Trials subject categories stopped accepting new papers.

### Preprint features

During the submission process authors upload either a complete article as a PDF file or a combination of Microsoft Word and figure files, which are then automatically converted into a single PDF file. Manually entered article information generates the HTML metadata that is viewable when the article first appears online (see below). Authors may also upload additional supplemental files, such as movies or supplementary figures and tables. DOIs are assigned after the authors have approved the PDF for posting. Article screening and posting typically takes 24–48 hours, barring any issues that need to be addressed by the authors before posting or occasional delays due to weekends or holiday periods. Papers initially post as author PDF files together with author-entered metadata and supplementary material. Full-text HTML and journal article tagging set (JATS) format XML generated by an external compositor are added 24– 48 hours later. These include in-line figures and linked references (Fig. 2) and are also used to generate a print friendly/PDF version with in-line figures.

**Fig. 2.**
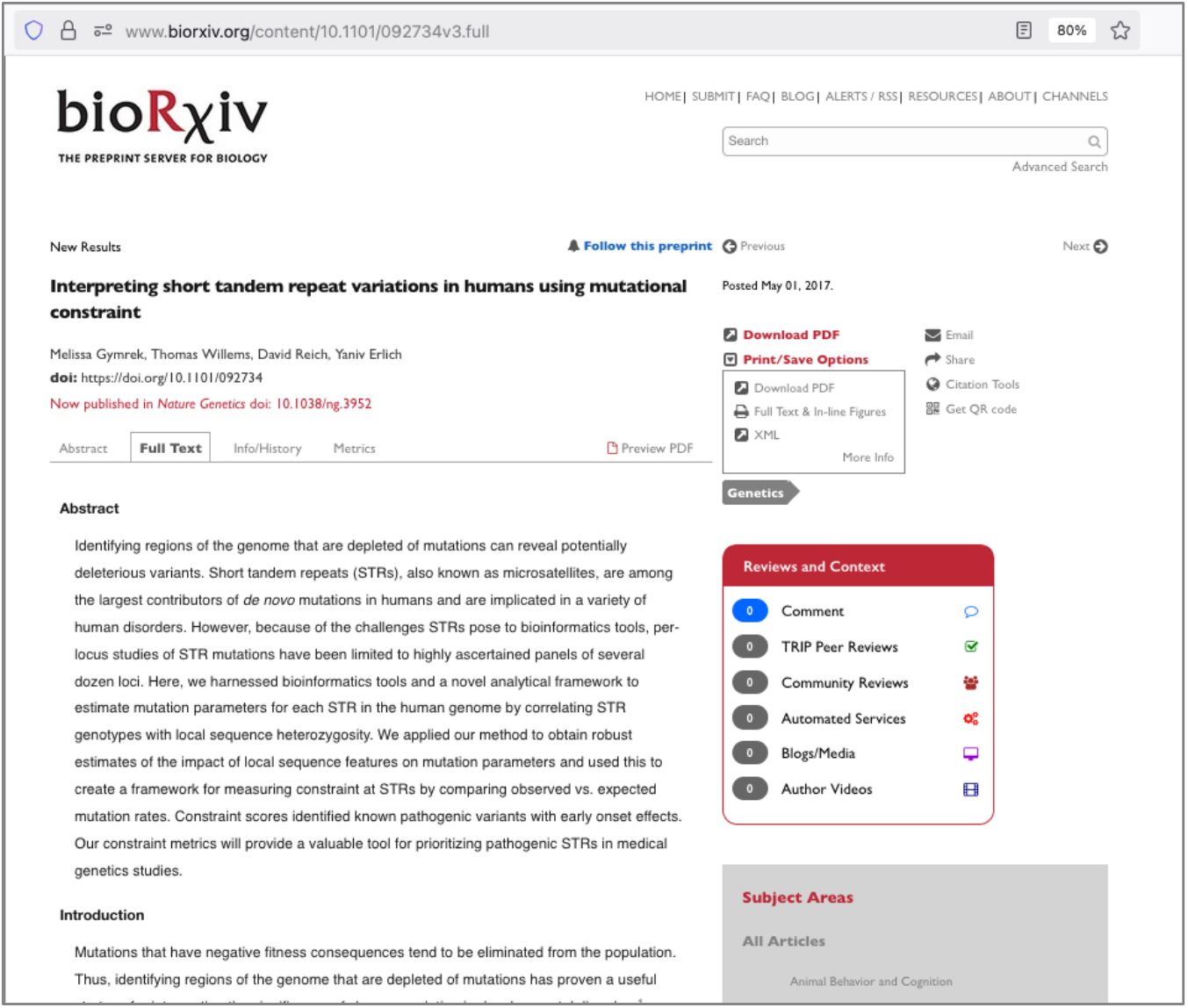
Screenshot showing a bioRxiv article HTML view. Content is displayed in full-text HTML, along with a download link for the authors’ PDF file, a print-friendly view with in-line figures, and a variety of additional information. The article history and links to earlier versions of the paper can be accessed via the Info/History tab.

Other elements displayed include the single subject category and article type (New Results, Contradictory Results or Confirmatory Results) selected by the authors, the article history (links to prior versions), and the authors’ choice of terms under which they wish to make the article available. These include various Creative Commons licenses, ‘all rights reserved’, CC0 Public Domain dedication, and a specific US government Public Domain option required for NIH employees. In addition to standard article metadata, authors may also provide ORCIDs and links to externally hosted data sets or code within a dedicated field. For revised versions of articles, they can also include a revision summary (version note) describing the changes they have made during revision. Additional elements viewable alongside articles include links to the final journal version of record when this appears, accepted article notifications for participating publishers, and article-level metrics.

### The bioRxiv dashboard

One of the goals of bioRxiv is to alert readers to additional context and discussion around preprints. The bioRxiv dashboard provides a mechanism for aggregating and linking comments, independent peer reviews, automated analyses, community discussion, and conference videos (Fig. 3). These include formal peer reviews from journals and independent peer review services that participate in the transparent review in preprints (TRiP) project, which bioRxiv displays alongside preprints by using Hypothesis annotation technology. A similar approach is used to show outputs of automated services such as Sciscore (which extracts research resource identifiers, RRIDs) and Sciencecast (which creates AI summaries to broaden accessibility of the material). Links are provided to community discussion sites and blog/media coverage of preprints, while online commenting is provided via the Disqus web plug-in.

**Fig. 3.**
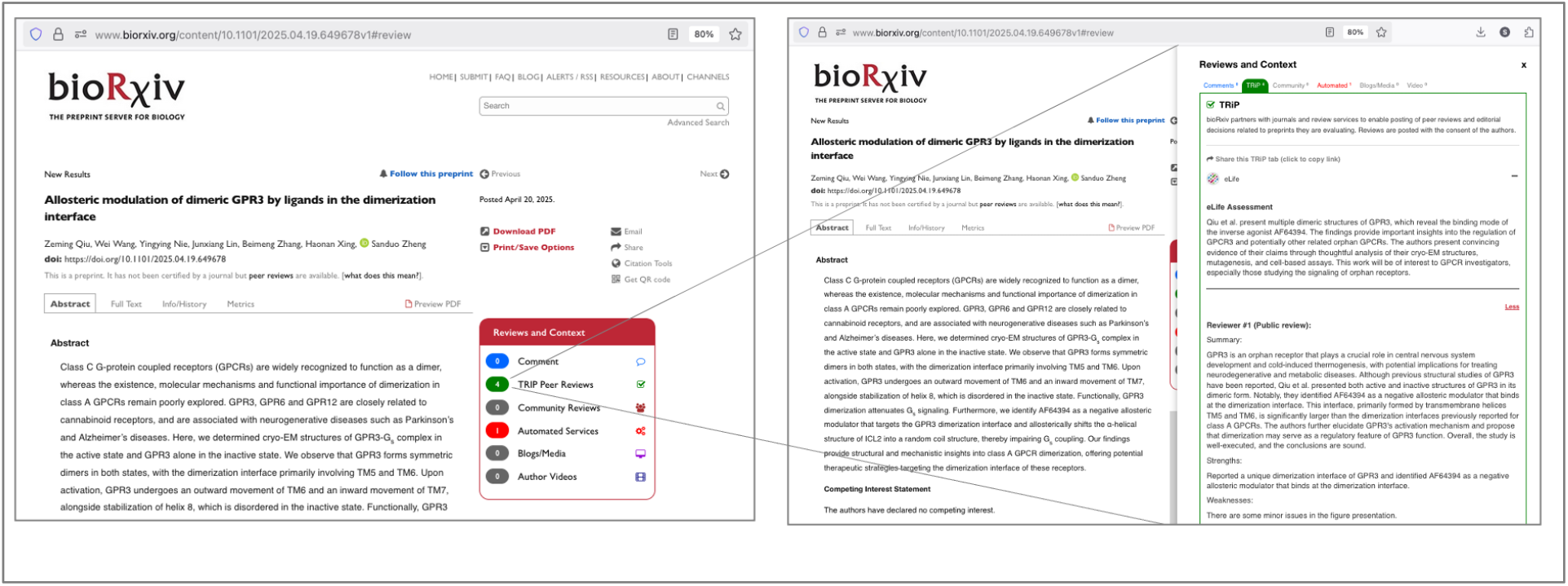
Screenshot showing the bioRxiv dashboard. The bioRxiv dashboard (bottom right, left panel) displays the number of comments, peer reviews, community discussion, automated services, blog/media posts, and author videos associated with a preprint. When the dashboard is opened (right panel) the commentary is shown in full or linked, depending on the nature of the content.

### Indexing and discovery

DOIs assigned to bioRxiv articles are deposited with the DOI-registration agency Crossref on the day of posting. Once the article is published in a journal, bioRxiv adds a link to the formally published version alongside the preprint and updates the Crossref DOI record with this information, which is subsequently available via bioRxiv (api.biorxiv.org) and Crossref (api.crossref.org) APIs. bioRxiv identifies preprint–journal article matches through a variety of scripts that search PubMed and Crossref databases for title and author matches. Matched authors are then alerted and have the opportunity to remove the link if the match is incorrect and/or supply matches for articles that have not been identified. bioRxiv extends this approach to articles that have been retracted from journals, so this information can also be displayed alongside relevant preprints.

bioRxiv includes numerous built-in search and alert features and is indexed by a variety of third-party discovery tools. Readers can browse the site by subject category or using the Solr-powered search feature within the hosting site. Personalized email alerts for specific search terms can also be generated, and subject-category-specific RSS feeds and social media accounts provide additional mechanisms for content alerts. Readers can also sign up to follow preprints and receive alerts when they are revised, commented on, or published in a journal. bioRxiv is indexed by generic search engines, as well as dedicated literature-discovery engines such as Google Scholar, Web of Science, and Europe PubMed Central. APIs and a dedicated text and data mining (TDM) repository are also available.

### Manuscript transfers and connections

To reduce the burden on authors who wish to submit to both bioRxiv and journals—and to further encourage preprint posting—we have developed bioRxiv-to-journal (B2J) and journal-to-bioRxiv (J2B) streams that allow authors to transfer articles between bioRxiv and journal submission systems. This means that authors need only upload files and manually enter core metadata once, saving them significant time and effort, although some journals require additional journal-specific metadata following B2J that must be entered separately at the journal submission site. B2J and J2B use the standard File Transfer Protocol (FTP) to transmit a ZIP archive containing XML metadata and manuscript files in a way that can easily be generated/ingested by journal submission systems. B2J and J2B pre-date, and in some ways inspired, the Manuscript Exchange Common Approach (MECA; Sack, 2018) to which B2J and J2B have been adapted.

The B2J and J2B manuscript transfer services are not just available to journals. B2J can be used to transfer papers from bioRxiv to journal-independent peer review services, such as Peer Community In (PCI) and Review Commons. Meanwhile, authors can also transfer manuscripts or metadata to signal their interest in third-party services like Dataseer, Dryad or q.e.d Science via a process termed B2X.

### Withdrawals

Manuscripts posted on bioRxiv receive DOIs and thus are citable and part of the scientific record. In addition, they are indexed by third-party services, creating a permanent digital presence independent of bioRxiv records. Consequently, bioRxiv’s policy is that papers cannot be removed, except in exceptional cases for legal reasons or where there might be serious risk of harm.

Authors can, however, have articles marked as “Withdrawn” if they no longer stand by the findings/conclusions or if they acknowledge fundamental errors in the article. In these cases, the default view becomes a withdrawal statement providing an explanation for the withdrawal, but the original article is still accessible via the article history tab. In rare cases, an article can be withdrawn by bioRxiv itself as a consequence of unethical behavior by an author or a technical error made by bioRxiv or its technology partners.

Withdrawn articles are clearly identified within the bioRxiv website. Ensuring that this signal is perpetuated within the ecosystem and that such withdrawals are effectively identified, indexed and displayed by third-party services is an area currently being investigated. For example, the word “Withdrawn” is added at the beginning of the title metadata to facilitate identification in Google Scholar.

### bioRxiv by the numbers

Below we summarize a series of data sets related to preprints posted on bioRxiv. The numbers are current at the time of writing, but we wish to alert readers to the fact that many of these metrics are updated in real time and available to interested readers at api.biorxiv.org.

Figure 4 shows the number of bioRxiv posts since 2013. Over this period submissions grew considerably from a handful to more than 4000 per month in 2025 (Fig. 4A). At the time of writing, the total number of first submissions to bioRxiv is more than 310,000. The proportion of manuscripts that are revised has remained fairly constant and is currently 26%. Most papers are revised only once (if at all) but some are revised multiple times (8% have two revisions; 4% have three or more revisions). 500 papers were withdrawn between 2018 and 2025. This represents a withdrawal rate of 0.17%, which compares to the ∼0.2% of journal articles that are retracted (Van Noorden, 2023).

**Fig. 4.**
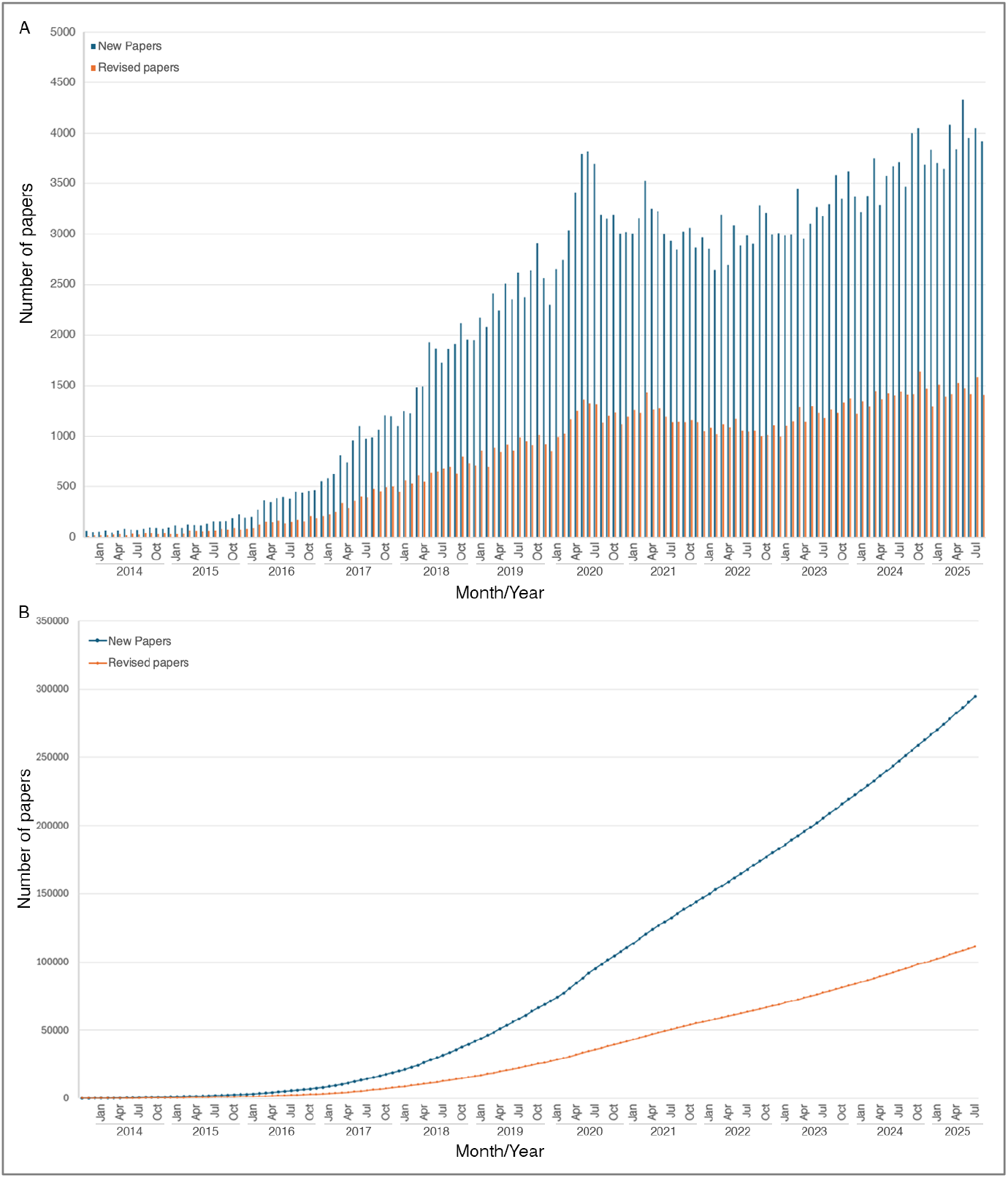
The growth of bioRxiv. A. Monthly submissions to bioRxiv. New articles are in blue; revised articles are in red. B. Total number of papers on bioRxiv. New articles are in blue; revised articles are in orange.

Table 1 shows the fractions of articles within different subject categories across a 13-year period. Initially bioRxiv was dominated by papers in genomics, bioinformatics and evolutionary biology, but the percentages contributed by other subdisciplines have increased, most notably in neuroscience (Table 1). This is consistent with the experience of arXiv, which was initially dominated by high-energy physics but subsequently began to attract papers from other disciplines in large numbers (Ginsparg, 2011). bioRxiv preprints have been deposited by authors from 209 different countries, the most common being the USA, UK, Germany, and China. The most prolific institutions are Stanford University, University of Oxford, and University of Cambridge (Table 2). The distribution of licenses chosen by authors is relatively constant with a small uptick in CC BY licensing following re-ordering of the selection options in 2025: 30% CC BY-NC-ND, 27% CC BY, 25% all rights reserved, 12% CC BY-NC, 4% CC BY-ND, 2% CC0/Public Domain.

**Table 1.** bioRxiv papers by subject category.

| Subject Category | 2013 (%) | 2014 (%) | 2015 (%) | 2016 (%) | 2017 (%) | 2018 (%) | 2019 (%) | 2020 (%) | 2021 (%) | 2022 (%) | 2023 (%) | 2024 (%) | 2025 (%) |
| --- | --- | --- | --- | --- | --- | --- | --- | --- | --- | --- | --- | --- | --- |
| Animal Behavior And Cognition* |  | 1.15 | 1.14 | 1.55 | 1.62 | 1.71 | 1.58 | 1.44 | 1.51 | 1.56 | 1.57 | 1.68 | 1.54 |
| Biochemistry | 0.92 | 0.57 | 1.14 | 2.09 | 2.78 | 3.08 | 3.28 | 4.3 | 4.14 | 4.46 | 4.53 | 4.6 | 4.59 |
| Bioengineering | 0.92 | 0.57 | 1.31 | 0.79 | 1.48 | 2.06 | 2.52 | 2.95 | 3.11 | 3.47 | 3.71 | 3.89 | 3.99 |
| Bioinformatics | 12.84 | 18.25 | 17.83 | 15.05 | 11.34 | 9.28 | 8.04 | 7.98 | 7.92 | 8.4 | 8.29 | 8.27 | 8.59 |
| Biophysics | 3.67 | 2.53 | 2.11 | 3.85 | 4.08 | 4.49 | 4.38 | 4.71 | 5.05 | 5.04 | 4.79 | 4.58 | 4.63 |
| Cancer Biology | 4.59 | 1.72 | 2.68 | 2.07 | 3.13 | 3.45 | 3.73 | 3.99 | 3.9 | 4.17 | 4.57 | 4.8 | 5.04 |
| Cell Biology | 1.83 | 2.07 | 1.14 | 3.47 | 4.74 | 5.06 | 5.36 | 5.86 | 5.83 | 6.11 | 6.29 | 6.02 | 5.92 |
| Clinical Trials**† |  |  | 0.06 | 0.04 | 0.17 | 0.23 | 0.13 |  |  |  |  |  |  |
| Developmental Biology | 2.75 | 2.53 | 1.77 | 1.87 | 2.93 | 3.38 | 3.02 | 2.99 | 3.22 | 3.13 | 3 | 3 | 2.75 |
| Ecology | 8.26 | 4.94 | 4.79 | 4.94 | 3.85 | 3.97 | 4.4 | 4.62 | 4.59 | 4.13 | 4.03 | 4.06 | 4.27 |
| Epidemiology***† |  |  |  | 1.66 | 2.85 | 2.95 | 1.86 |  |  |  |  |  |  |
| Evolutionary Biology | 15.6 | 20.21 | 18.46 | 11.16 | 7.44 | 5.59 | 5.14 | 4.58 | 4.55 | 4.5 | 4.25 | 3.97 | 3.81 |
| Genetics | 8.26 | 8.15 | 10.71 | 7.73 | 6.69 | 5.08 | 4.8 | 3.56 | 3.14 | 3.09 | 2.75 | 2.55 | 2.43 |
| Genomics | 11.01 | 15.04 | 13.45 | 10.75 | 7.61 | 6.19 | 5.41 | 5.01 | 4.56 | 4.22 | 4.06 | 4.08 | 4.13 |
| Immunology | 1.83 | 1.49 | 0.8 | 1.09 | 1.57 | 2.66 | 3.25 | 4.41 | 5.11 | 4.71 | 4.34 | 4.44 | 4.62 |
| Microbiology | 2.75 | 1.61 | 3.3 | 5.54 | 6.39 | 8.95 | 9.02 | 10.63 | 10.16 | 10.04 | 9.38 | 9.57 | 9.43 |
| Molecular Biology | 1.83 | 1.61 | 1.14 | 1.96 | 2.58 | 3.44 | 3.69 | 3.93 | 4.03 | 4.1 | 4.42 | 4.33 | 4.3 |
| Neuroscience | 8.26 | 7.12 | 8.26 | 14.9 | 18.6 | 18.29 | 18.99 | 17.53 | 17.37 | 17.63 | 18.28 | 18.34 | 18.36 |
| Paleontology | 0 | 0.11 | 0.17 | 0.09 | 0.13 | 0.17 | 0.14 | 0.11 | 0.16 | 0.17 | 0.14 | 0.14 | 0.15 |
| Pathology | 0 | 0.11 | 0.28 | 0.28 | 0.3 | 0.55 | 0.65 | 0.7 | 0.71 | 0.63 | 0.75 | 0.74 | 0.78 |
| Pharmacology And Toxicology* |  | 0.23 | 0.28 | 0.45 | 0.73 | 0.8 | 1.02 | 1.29 | 1.25 | 1.15 | 1.21 | 1.26 | 1.27 |
| Physiology | 0 | 0.69 | 0.51 | 0.51 | 0.95 | 1.18 | 1.66 | 1.74 | 1.83 | 1.68 | 1.94 | 2.13 | 2.08 |
| Plant Biology | 4.59 | 1.26 | 1.94 | 2.26 | 2.75 | 2.63 | 3.39 | 3.69 | 3.88 | 3.71 | 3.67 | 3.86 | 3.63 |
| Scientific Comm. & Education | 0 | 0.57 | 0.46 | 0.75 | 0.6 | 0.7 | 0.77 | 0.6 | 0.47 | 0.5 | 0.36 | 0.29 | 0.26 |
| Synthetic Biology | 4.59 | 1.95 | 1.42 | 1.09 | 1.08 | 0.88 | 0.87 | 0.82 | 0.94 | 0.92 | 1.1 | 1.11 | 1.06 |
| Systems Biology | 3.67 | 4.71 | 4.16 | 3.7 | 3.19 | 2.79 | 2.31 | 2.06 | 2.01 | 1.96 | 1.99 | 1.72 | 1.81 |
| Zoology | 1.83 | 0.8 | 0.68 | 0.38 | 0.42 | 0.46 | 0.59 | 0.53 | 0.57 | 0.53 | 0.6 | 0.55 | 0.54 |

\*introduced 2014 \*\*introduced 2015 \*\*\*introduced 2016 †closed 2020

**Table 2.** Top-ten preprinting institutions.

| Institution | Country | Preprints |
| --- | --- | --- |
| Stanford University | USA | 2739 |
| University of Oxford | UK | 2144 |
| University of Cambridge | UK | 2003 |
| University of Washington | USA | 1714 |
| University of Pennsylvania | USA | 1662 |
| University of Michigan | USA | 1632 |
| Columbia University | USA | 1268 |
| University College London | UK | 1255 |
| Johns Hopkins University | USA | 1239 |
| Yale University | USA | 1222 |

bioRxiv usage grew significantly in the first six years (Sever et al., 2019a) but has stabilized since the pandemic. The site currently receives >7 million abstract views per month and ∼4 million PDF downloads per month. The numbers for full-text HTML views are currently around half the level of PDF downloads, but full-text HTML was introduced only in 2019 and is unavailable until 24–48 hours after PDF posting, so immediate feeds/alerts will favor the PDF file.

Most bioRxiv preprints are ultimately published in traditional journals. Matching algorithms find that ∼80% of bioRxiv preprints are published by journals after a 3-year period sufficient for most papers to have passed through review and revision cycles to acceptance. This fraction is consistent with findings for arXiv (Larivière et al. 2013). When a preprint is published in a journal, a prominent link to the publication is inserted above its abstract. Such a link may be absent because the title and/or authorship of the manuscript have changed sufficiently during publication to make it no longer identifiable by matching algorithms or because the paper is still under consideration at a journal. An 80% publication rate may therefore be an underestimate.

Articles that first appeared as preprints on bioRxiv have now been published in more than 5000 journals. Supplementary Table 1 shows the number of preprints for the 10-most-common destination journals at the time of writing. Comprehensive, updated numbers are available at api.biorxiv.org. The journals that publish bioRxiv preprints represent a wide spectrum of specialties and are both open access and subscription-based. Unsurprisingly, the mega-journals *PLOS ONE* and *Scientific Reports* are highly represented. Journals such as *eLife* that participate in both B2J and J2B also receive significant numbers of papers. Journals that cover subdisciplines highly represented in bioRxiv are more likely to receive relatively high numbers of papers compared with equivalent titles in less well-represented subdisciplines. The interval between the posting of a preprint and its publication in a journal is influenced by variables such as time to first submission, the number of serial submissions before acceptance, and the extent of revisions required by peer review. For all manuscripts on bioRxiv, the interval between availability on bioRxiv and journal publication currently averages 256 days (median 204 days).

One aspiration for preprints has been that they provide a mechanism for the community to provide feedback on papers for authors and readers. bioRxiv therefore includes an on-site commenting mechanism. It also displays peer reviews from journals and other peer review services and aggregates links to discussions elsewhere on the Web and in social media. 4% of papers include peer reviews as part of the TRiP partnership with journals and peer review services. 4% of papers currently display onsite comments, while just over 2% receive community reviews on third-party sites. There are extensive discussions of articles on social media, and authors receive private feedback via email (see below); so this may simply reflect the fact that on-site commenting is not the preferred medium for providing feedback. Additional cultural change is required for public commenting to become the norm.

### bioRxiv in the pandemic

bioRxiv and medRxiv became crucial resources for rapid sharing of research on COVID during the pandemic. The servers tagged COVID-related papers and provided a dedicated COVID collection that at the time of writing includes >30,000 papers. The first preprint on the novel coronavirus that came to be termed SARS-CoV-2 was posted on bioRxiv on January 19^th^, 2020 (Chen et al., 2020). Numerous important papers describing the structure, biology, and evolution of SARS-CoV-2 subsequently appeared first as preprints on bioRxiv. COVID submissions peaked in May 2020 (Fig. 5), when >2000 papers were received (bioRxiv and medRxiv). The number of COVID submissions declined through 2024, leveling off at ∼100/month in 2025. The challenges of rapid dissemination in a constantly developing area with important implications for public health underscored the importance of screening and responsible stewardship of preprints, in particular concerns about information that could present a public health risk (Sever et al., 2021).

**Fig. 5.**
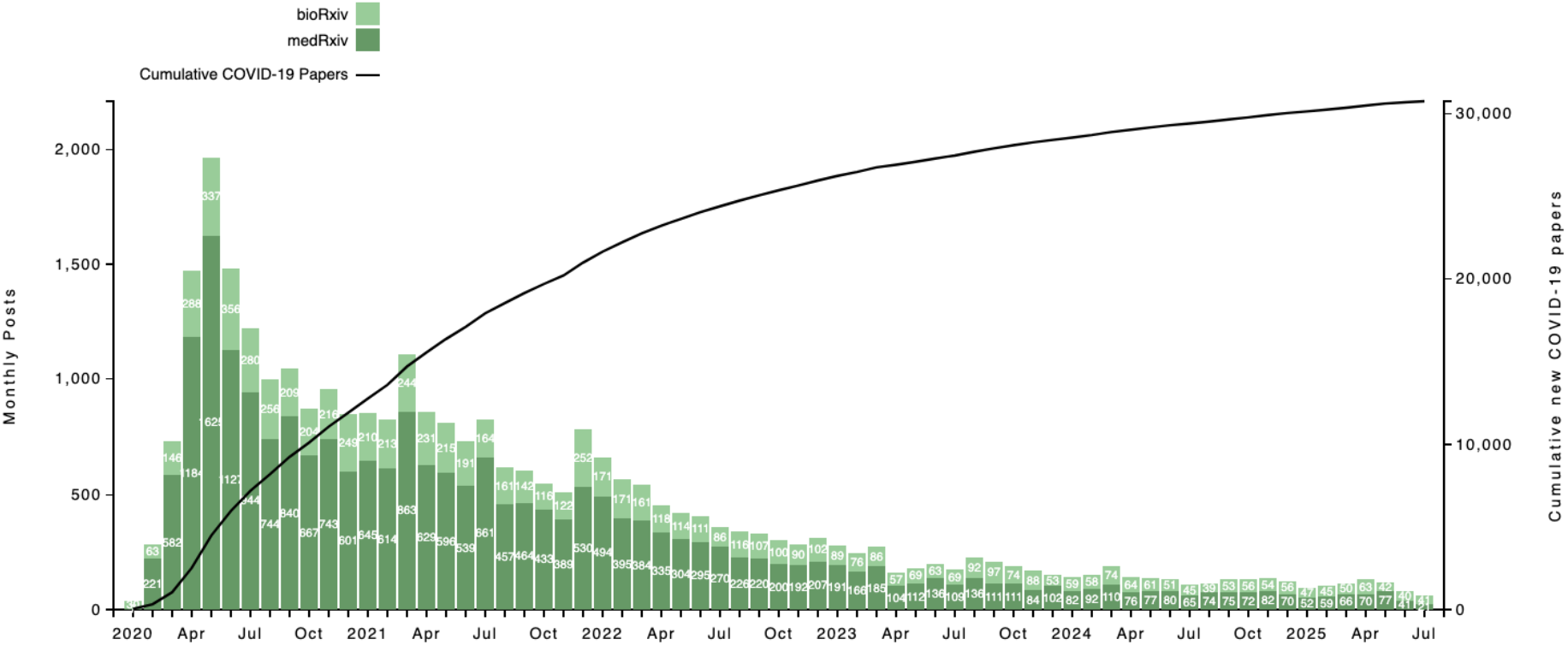
COVID papers on bioRxiv and medRxiv. Monthly posts of bioRxiv (light green bars) and medRxiv (dark green bars) COVID preprints are shown relative to the left-hand y-axis. Total COVID papers (black line) are shown relative to the right-hand y-axis.

### The 2023 bioRxiv survey

The first bioRxiv survey of >4000 bioRxiv users was conducted in 2017 (Sever et al., 2019a). We recently conducted a new survey of >7000 bioRxiv and medRxiv users in an effort to understand further how preprints are being used among life scientists and health researchers (bioRxiv and medRxiv, 2024). There will inevitably have been some self-selection bias in survey respondents and geographic representation. One should therefore be cautious about generalizing from the results. We nevertheless feel the results are informative and we highlight some of the key findings below.

bioRxiv uses a submission system in which authors can submit Microsoft Word documents and individual figure files and/or PDF files. This was based on the assumption that most authors in life sciences use Word to compose documents and contrasts with the submission process at arXiv, which focuses on LaTeX users. Figure 6 shows that 85% of bioRxiv survey respondents indeed use Word. A significant minority use LaTeX (22%) and it is important to emphasize that LaTeX users can submit to bioRxiv; they need simply create a PDF version of their paper as well. 44% of survey respondents use Google docs, but note that these authoring tools are not mutually exclusive. 10% of respondents state that they use large language models (LLMs). The survey was conducted in 2023, however, and so it is likely this fraction has increased. A variety of reference managers are used, including Zotero (32%), EndNote (28%), Mendeley (18%), and Papers (6%).

**Fig. 6.**
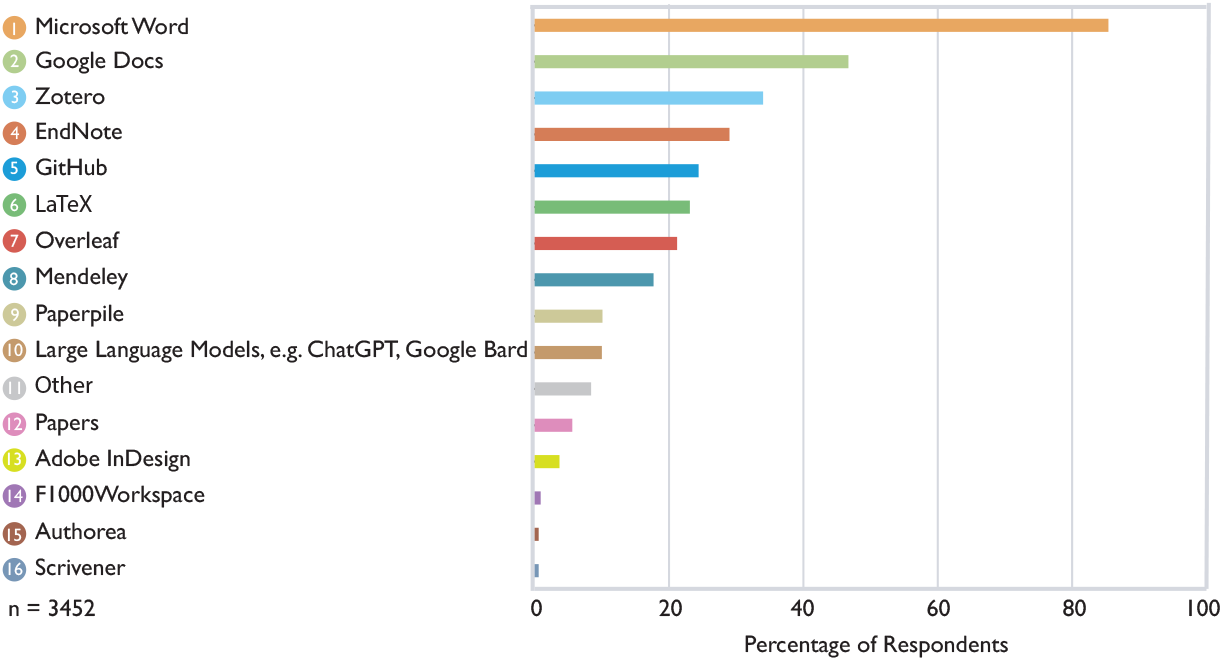
Authoring software used by authors. In a survey of bioRxiv users, authors were asked what software they use to prepare manuscripts.

There is much discussion among scientists, publishers and IT professionals about authoring technologies and a desire is often voiced for the development and adoption of new tools. The survey findings remind us that it is important to cater for the tools that people are already using, as well as new approaches, particularly when trying to incentivize adoption of new cultural practices such as preprint posting.

We also surveyed authors on their motivation for posting preprints and the consequences of posting. The survey revealed a variety of motivations for posting (Fig. 7), including increasing awareness of research (82%), staking a priority claim (56%), controlling when research is available (54%), a wish to cite work in a grant (48%) or job application (30%), and a desire to get feedback (38%). Most respondents (64%) also felt that immediate sharing of new results benefits the scientific enterprise.

**Fig. 7.**
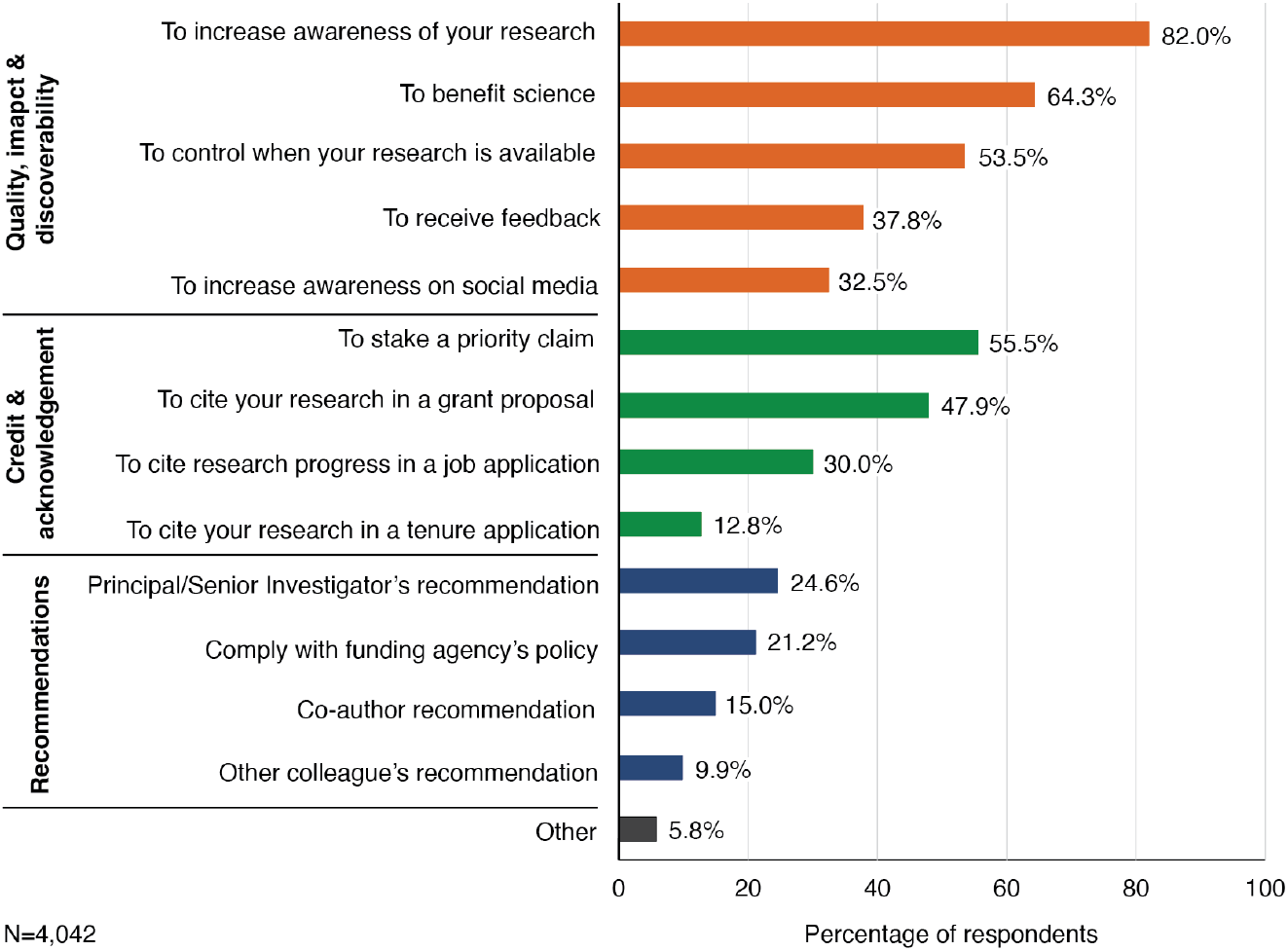
Motivations for posting work on bioRxiv. In a survey of bioRxiv users, authors were asked why they post manuscripts on the server.

Given the contrast between the anticipated desire for feedback and the relatively low volume of on-site commenting on bioRxiv, we were keen to learn more about the feedback authors received via different channels (Fig. 8). Importantly, 36% of authors said they had received feedback on preprints by email and 34% through conversations, neither of which bioRxiv can quantify directly. A further 32% had received feedback via Twitter/X and 9% had received feedback via bioRxiv’s online commenting section, a figure that indicates some sampling bias among survey respondents given the 4% figure for commenting noted above. Nevertheless, since 46% of surveyed authors express a strong desire for feedback via online comments, that desire is only partly being satisfied (Fig. 8). Perhaps this is because the technological solutions available are not ideal, but a more likely cause is the absence of meaningful rewards for commenting and providing online feedback within a community already pressed for time. Indeed, 58% of survey respondents had never provided feedback on a preprint. However, since the overwhelming majority of survey respondents indicated a wish to receive feedback via email (Fig. 8), there may be a balance to be struck between private and public channels.

**Fig. 8.**
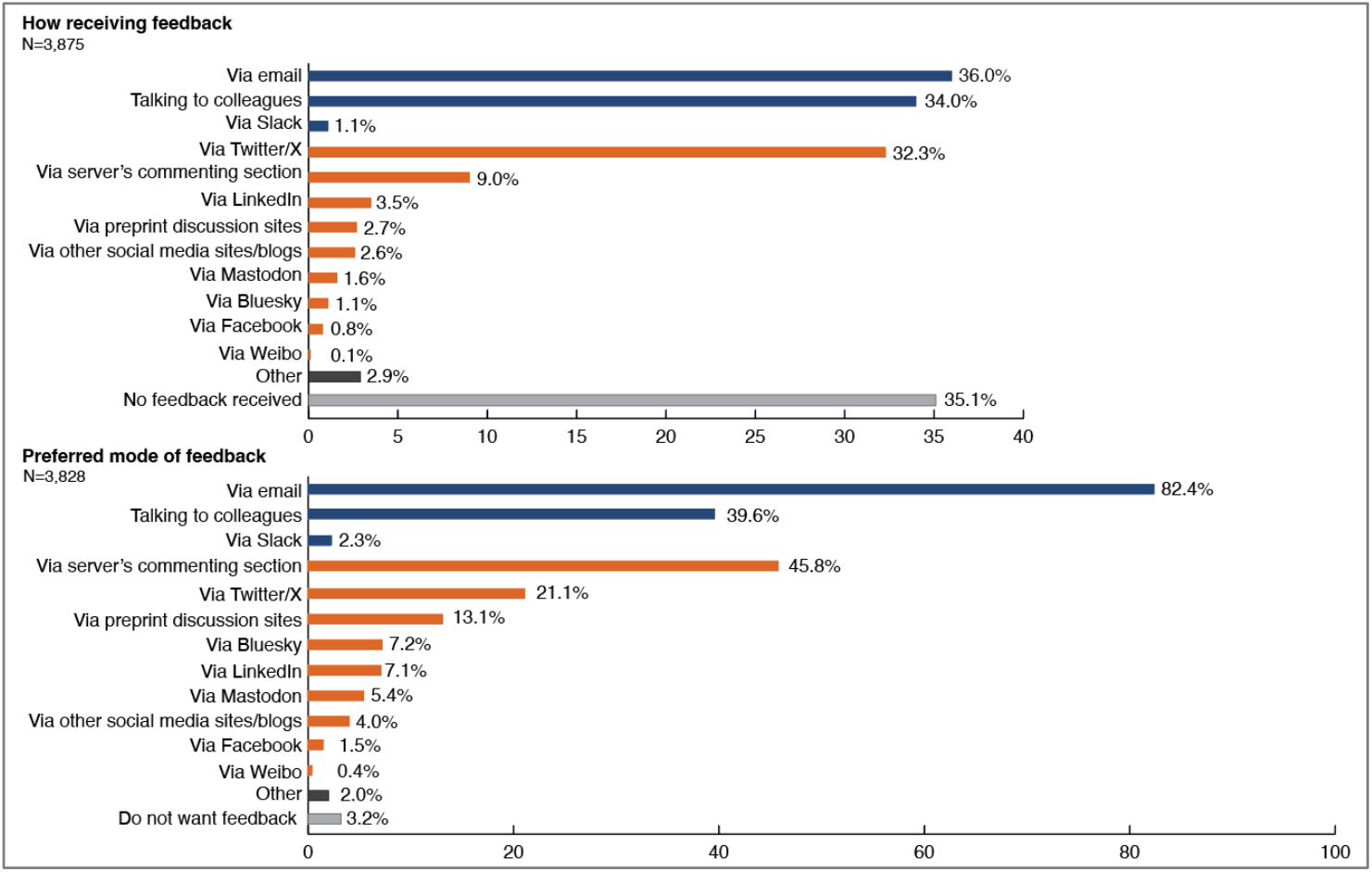
Feedback mechanisms. In a survey of bioRxiv users, authors were asked the mechanisms by which they have received feedback on papers posted on bioRxiv (upper panel) and how they would like to receive feedback on papers (lower panel).

Since motivations for preprint posting include both the desire to get work out early and the hope of receiving feedback, we asked when authors post preprints in the course of preparing submissions to journals. 30% of authors said they post weeks to months before they submit to their first-choice journal (17.6% post 1-4 weeks before; 12.5% post >1 month before); 55% of authors said they post a preprint the same week they submit to their first-choice journal (see Supplementary Table 2). These differences may reflect differences in whether authors wish to obtain feedback before submitting to journal.

Survey respondents reported that posting a preprint had helped in a variety of ways (Fig. 9). 78% said that it had increased awareness of their research. Others found that it had helped them meet people in their field (16%) and/or make progress in it (36%). A smaller number said that it had helped them get a job, grant or seminar invitation (8%, 13%, and 12%, respectively). 34% believe it helped them stake a priority claim, a major motivation for posting in the physical sciences (Ginsparg, 2011). The vast majority of authors (89%) had experienced no negative consequences of preprint posting. Only 2% believed that it had prevented them publishing in a specific journal by giving a competing group an advantage (bioRxiv and medRxiv. 2024). 4% felt it had limited their choice of journal, presumably because a small number of journals will not consider manuscripts previously posted on a preprint server.

**Fig. 9.**
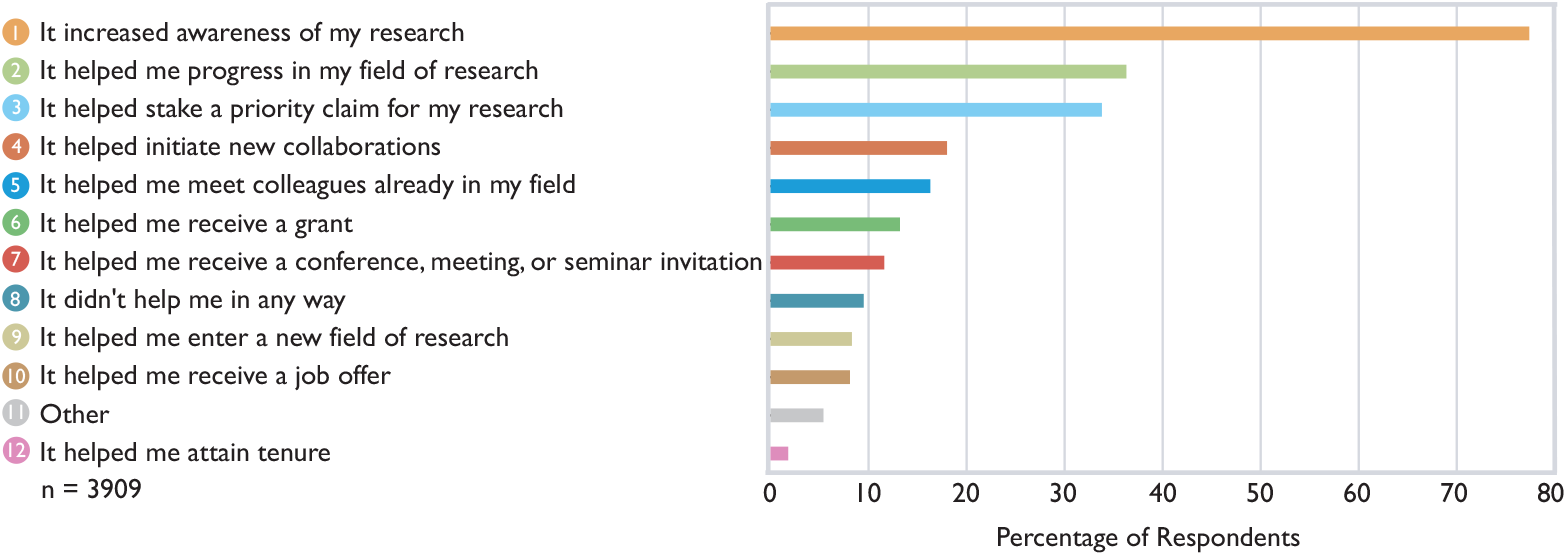
How preprints help authors. In a survey of bioRxiv users, users were asked how posting an article on bioRxiv has helped them in their careers.

## Discussion

bioRxiv has grown hugely in popularity since its launch in 2013, reflecting an increasing desire within the life science community for rapid and open dissemination of results. There is a positive feedback loop operating, with greater usage and increased familiarity with bioRxiv driving further adoption of the practice of preprinting and its spread to new subdisciplines. The growth of bioRxiv has also helped prompt the launch of numerous similar servers in other fields (e.g. chemRxiv, SocArXiv, PsyArXiv and EarthArXiv) and inspired the creation of medRxiv.

A number of other factors have contributed to preprint adoption. These include changes in many publishers’ policies allowing their journals to consider papers previously posted to preprint servers. ^1^ Furthermore, journals such as the Public Library of Science (PLOS) titles and *eLife* now actively encourage or require preprint posting by authors (PLOS, 2018). Similarly, many funders now allow or encourage inclusion of preprints in grant applications and even mandate posting in some cases (e.g. Aligning Science Across Parkinson’s, CZI, the Gates Foundation, Howard Hughes Medical Institute, and the Michael J. Fox Foundation), and the NIH recognizes preprints as interim research products (NIH, 2017; NIH, 2019). Meanwhile, the career benefits to authors that preprints afford are increasingly clear to both early career researchers and senior investigators (Hindle and Sever, 2024).

It is important to stress, however, the extent to which the research community itself has been the driver of preprint adoption. Genetics and genomics researchers were particularly early adopters and vocal advocates of bioRxiv, and awareness of bioRxiv spread fast among the bioinformatics and evolutionary biology communities. More formal initiatives such as ASAPbio followed and helped spread the word within other subdisciplines, in particular cell and developmental biology. The establishment of preprint discussion sites such as preLights (Brown and Pourquié, 2018) and others has also contributed. bioRxiv has benefited enormously from the enthusiasm with which individual scientists in research communities worldwide have embraced preprints and become active advocates for this approach to dissemination (Sever, 2026). Social media have played an important part in spreading preprint awareness among scientists, alerting readers to individual articles, and providing a conduit for automated article feeds that bioRxiv provides.

It will be interesting to see how greater adoption of preprints further stimulates the evolution of scientific communication and peer review in particular. Anecdotal evidence indicated from the beginning that journal editors were soliciting papers from authors who post preprints on bioRxiv and journals such as *PLOS Genetics* and *Proceedings of the Royal Society B* appointed editors specifically tasked with such recruitment. B2J is also making the process of journal submission easier for bioRxiv authors. The additional scrutiny of papers from the community prior to journal submission/publication has the potential to improve the quality of papers and optimize peer review. As dissemination and evaluation become decoupled, the pressure to evaluate quickly may be relieved, reducing errors and allowing more thorough and potentially tailored peer review. The very existence of preprints is promoting experimentation with the peer review process at journals (Brainard, 2019) and elsewhere. This is particularly timely given ongoing discussions about the potential for more open and/or transparent peer-review processes (ASAPbio, 2018), additional trust signals (Hall Jamieson et al., 2019), and portable peer review (EMBO and ASAPbio, 2019). Going forward bioRxiv will continue to facilitate such new initiatives, as it has journal transfers and linking, community discussion, and reproducibility efforts.

bioRxiv also intends to take advantage of advances in technology and changes in tools used by life scientists. While plagiarism checks are already largely automated, scientific screening is currently performed by individuals. It is unlikely that human judgment could be entirely replaced, but AI approaches offer the hope of automated processes that augment and facilitate human screening. The submission process includes automated aspects of file processing such as PDF generation and verification but still requires manual data entry for other aspects. Improvements in automated text extraction and tagging could make this more efficient, as could a new generation of authoring tools that allow easier generation of XML/HTML. The format of scientific articles has changed little over the years — in many respects it remains tied to a layout dictated by the requirements of print journals. However, the variety of file types employed for different data types, use of tools such as Jupyter notebooks, and broader recognition of code as an integral part of scientific methods and results mean that the content encompassed by the term “research paper” will change, and so too will the outputs with the increasingly anachronistic description ‘preprint’.

## Concluding remarks

Physicists, computational scientists and mathematicians have been sharing research papers prior to peer review and formal publication for over three decades. bioRxiv has made this practice widespread in the life sciences and inspired preprint servers in many other disciplines. The decoupling of dissemination and evaluation combined with rapid online posting accelerates awareness of new work and so can increase the pace of research itself. Preprints provide a route to the long-desired goal of making research information freely and immediately available to anyone (Sever et al., 2019b). They also create opportunities for evolution of the publishing ecosystem. Broad adoption of preprints, together with technological advances, has the potential to create a more open, equitable and efficient system for the distribution, assessment and archiving of scholarly information.

## Acknowledgments

We would like to thank all of those who have worked on and advocated on behalf of bioRxiv, in particular CSHL colleagues Dinar Yunusov, Joanne McFadden, Inez Sialiano, Mary Mulligan, Tara Kulesa, Justin Kinney, Bruce Stillman, Terri Grodzicker, Hillary Sussman, Laura DeMare, Dorothy Oddo, Kathy Bubbeo, Denise Weiss, Robert Redmond, Katherine Kelly, and Carol Brown and the medRxiv Co-Founders Theo Bloom, Harlan Krumholz, Claire Rawlinson and Joseph Ross. Thanks also to Robert Lourie, Jeremy Freeman, Cori Bargmann, Scott Fraser, Dario Tarborelli, Carly Strasser, Anurag Acharya, Fiona Watt, Leslie Vosshall, Graham Coop, Daniel MacArthur, Leonid Kruglyak, Prachee Avasthi, Jessica Polka, Joseph Pickerel, Yaniv Erlich, Steve Shea, Jessica Tollkuhn, Richard Murray, Chris Gunter, Casey Greene, Michele Avissar-Whiting, Bodo Stern, Michael Hoffman, Jim Woodgett, Michael Eisen, Veronique Kiermer, Allison Mudditt, Louise Page, Thomas Lemberger, Bernd Pulverer, Tracey DePellegrin, Eric Topol and Katherine Brown for advice and support.

This document was created using an adapted Word preprint template developed by the Finkelstein lab (Finkelstein, 2018).

## Data availability

Data underlying the results presented here are available at api.biorxiv.org. The full data set (minus any identifying information) for the bioRxiv author survey is provided in the reference supplied (bioRxiv and medRxiv. 2024).

## Ethics statement

The research in this study was reviewed and approved by the Cold Spring Harbor Laboratory IRB (2118552-1) and deemed exempt under 45 CFR 46.104 (d) 2.

## Supplementary Material — Survey design, execution and analysis

The 2023 survey was generated using Survey Monkey and comprised 39 multiple-choice questions and 8 open-ended questions. Questions were divided across user type; authors, readers, and non-users viewed a maximum of 42, 25, and 4 questions, respectively.

To understand how preprints have impacted our users, we focused the survey on bioRxiv/medRxiv authors and readers. To increase awareness of the survey, we added a banner to the bioRxiv/medRxiv homepages and all preprint webpages and to the submission system. We also emailed bioRxiv/medRxiv authors. The survey was completed by 7,135 individuals during the period it was open (October 25 to December 19, 2023).

The survey data were downloaded from SurveyMonkey and parsed using the R package “surveymonkey”, available from https://github.com/mattroumaya/surveymonkey, and then converted to long format. To avoid survey responses being used to identify individuals, free-text answers and respondent identifiers were removed. This dataframe was further processed using Rstudio (Posit Team, 2025) with R version 4.5.1. Briefly, the first question (“How do you use bioRxiv and/or medRxiv (Select all that apply)?”) was used to label users as ‘author’, ‘reader’ (i.e. non-author user) or ‘non-user’. Some respondents selected the option “I do not use bioRxiv or medRxiv” and another one; they were disqualified (labeled ‘non-user’) for the purposes of our analysis. They were also labeled as bioRxiv or medRxiv users based on that same question. Only survey data from bioRxiv users are used in the analysis presented in this preprint. Graphs with the percentages of respondents who selected each option were generated in RStudio using the ggplot package (Wickham, 2016) and modified using Adobe Illustrator 2025.

A de-identified version of the SurveyMonkey data in wide format is available in Zenodo (bioRxiv and medRxiv. 2024).

**Supplementary Table 1.** Top 10 journal destinations for bioRxiv preprints (2025)

| **Journal** | **Papers** | **Share** |
| --- | --- | --- |
| *Nature Communications* | 993 | 5.75% |
| *eLife* | 835 | 4.83% |
| *Proceedings of the National Academy of Sciences* | 410 | 2.37% |
| *PLOS ONE* | 380 | 2.2% |
| *Scientific Reports* | 324 | 1.88% |
| *Cell Reports* | 275 | 1.59% |
| *Science Advances* | 255 | 1.48% |
| *PLOS Computational Biology* | 239 | 1.38% |
| *Nucleic Acids Research* | 224 | 1.3% |
| *Communications Biology* | 205 | 1.19% |

**Supplementary Table 2.**
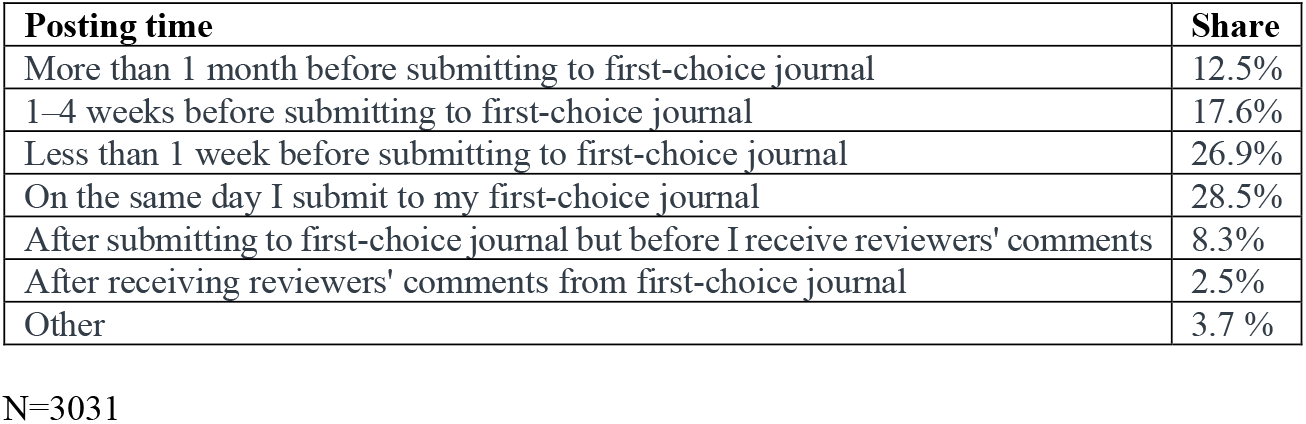
Preprint posting times reported by survey respondents.

| **Posting time** | **Share** |
| --- | --- |
| More than 1 month before submitting to first-choice journal | 12.5% |
| 1–4 weeks before submitting to first-choice journal | 17.6% |
| Less than 1 week before submitting to first-choice journal | 26.9% |
| On the same day I submit to my first-choice journal | 28.5% |
| After submitting to first-choice journal but before I receive reviewers' comments | 8.3% |
| After receiving reviewers' comments from first-choice journal | 2.5% |
| Other | 3.7 % |

N=3031

## Footnotes

1 Compare the current Wikipedia page listing academic journal policies (https://en.wikipedia.org/wiki/List_of_academic_journals_by_preprint_policy) with earlier versions of this page (e.g. https://web.archive.org/web/20130604021231/https://en.wikipedia.org/wiki/List_of_academic_journals_by_preprint_policy)

